# In pemphigus, cell detachment, but not autoantibody binding, induces cell-wide, long-lasting transcriptomic and proteomic changes

**DOI:** 10.1101/2025.02.10.637416

**Authors:** Veronika Hartmann, Sen Guo, Siavash Rahimi, Uta Radine, Danielle Malheiros, Amanda Salviano-Silva, Valeria Bumiller-Bini Hoch, Gabriel A. Cipolla, Veronica Calonga-Solis, Axel Künstner, Imke Burmester, Ralf J. Ludwig, William V.J. Hariton, Christoph M. Hammers, Eliane J. Müller, Hauke Busch, Jennifer E. Hundt

## Abstract

Desmoglein 1 (DSG1) and desmoglein 3 (DSG3) are adhesion molecules that maintain intercellular connections between epidermal, hair follicle, and mucosal keratinocytes. Autoantibodies (AAbs) targeting these molecules ultimately lead to the blister formation characteristic of pemphigus vulgaris (PV) or pemphigus foliaceus (PF). To investigate the molecular events following autoantibody binding up to 48 hours, we quantified transcriptome and proteome dynamics during split formation in a human skin organ culture (HSOC) model for PV and in a 2D cell-culture model for PV and endemic PF. Treatment of the cells in 2D culture with PX43, a single-chain variable fragment targeting DSG1/3, or with endemic PF anti-DSG1 IgG yielded neither a significant transcriptome nor a proteome response over time relative to the respective IgG controls. When treating the HSOC model with mouse antibody AK23 (targeting DSG3) or with PX43, only the latter induced split formation. In the absence of split formation, no differentially regulated pathways were detected at the transcriptomic level. Split formation, observed as early as 5 hours post-injection, was associated with significant and sustained upregulation of IFNγ and TNFα-related genes, mediated by upstream NFκB, MAPK, and JAK-STAT pathways. The gene expression changes, corroborated by proteomics data, were strongly correlated with early wounding and keratinocyte detachment, as well as the transcriptome profile in the skin from PV patients, while inversely associated with keratinocyte differentiation and cell stretching. The co-occurrence of well-wide and long-lasting transcriptome and proteome responses with split formation suggests that PF-IgG and PV-related AAbs neither induce downstream transcriptome nor proteome changes directly. Rather, these changes appear to be secondary effects resulting from reduced adhesion and mechanical-stress-induced split formation of keratinocytes *in vitro* and most likely in patient skin *in vivo*.

## Introduction

Pemphigus diseases are a group of rare, life-threatening autoimmune blistering disorders characterized by the loss of keratinocyte adhesion (acantholysis) in the epidermis and mucous membranes (1,2). This loss of adhesion is primarily caused by pathogenic autoantibodies (IgG) targeting desmosomal adhesion proteins, including desmoglein 1 (DSG1) and desmoglein 3 (DSG3). Pemphigus vulgaris (PV) and pemphigus foliaceus (PF), the two major subtypes, differ in their autoantibody profiles and clinical presentation. While PV targets DSG3 and DSG1, leading to mucocutaneous blistering, PF predominantly involves DSG1, causing superficial skin blistering.

While the precise triggers for AAb production in pemphigus are hitherto not completely understood, genetic predisposition (3–6) and environmental factors, such as infections, medications, and ultraviolet radiation (7–10), are believed to contribute to the initiation of autoimmunity. Treatment of PV and PF involves a combination of systemic medications to control disease activity, reduce blister formation, and promote healing (8). These include immunosuppressive agents, systemic and topical corticosteroids, or biological therapies such as rituximab to target B cells, which play a role in the production of autoantibodies (11). Because of the obvious treatment gap between immediately effective corticosteroids, having many side effects, and the considerable time for rituximab to take effect, research in pemphigus additionally focuses on elucidating and targeting cellular pathways involved after binding of PV AAbs to the epidermal keratinocytes (12,13).

PV is considered a desmosome turnover disorder (14–17). Desmosomes are stable yet dynamic structures for tissue adherence, reorganization, and wound healing (18). Their constituent proteins are highly regulated by posttranslational modifications governing function within the desmosome (19,20). Desmosome dynamics are intricately woven into a vast pathway and protein interaction network (21). PV AAb binding has been shown to reduce the numbers and size of desmosomes, reducing the overall adherence strength of neighboring cells (22,23). Still, a longstanding debate persists regarding the role of PV AAbs, i.e., whether they actively initiate cellular signaling, alter the keratinocyte phenotype, or simply act as passive witnesses in disease progression by remodeling and weakening desmosomes and cell adhesion (24,25).

Three non-exclusive mechanisms are currently under discussion: (i) steric hindrance of AAbs via direct interference with DSG1 and DSG3 molecules (25), (ii) internalization and depletion of desmogleins 1 and 3 from the cell membrane, changing desmosome homeostasis (14,18), and (iii) activation of downstream signaling events (9,12). Treatment of keratinocytes with PV IgG has shown transient, sequential phosphorylation of SRC (26,27), epidermal growth factor receptor (EGFR) (22,28,29), endoplasmatic reticulum stress (30), and p38 mitogen-activated protein kinase (MAPK), starting within minutes and lasting up to 6 hours and indicating potential downstream effects (26). Phosphorylation correlates with keratinocyte shrinkage, actin depolymerization, and keratin aggregation (26). However, whether this phosphorylation is a cause of the AAbs, or a consequence of disturbed desmosome signaling and weakened cell-cell attachment remains unclear. Modulating signaling pathways controlling desmosome internalization, stability, and turnover (12) have long been studied concerning abrogating PV AAb-induced acantholysis *in vitro*. Inhibiting p38 MAPK, EGFR, Rho GTPase, MYC proto-oncogene, or Ca^2+^ influx pathways 1-2 hours before AAb treatment were shown to reduce fragmentation in a 2D keratinocyte dissociation assay (9,31) or split formation in a 3D human skin organ culture (HSOC) model for PV (12). Over 30 different molecular targets have been reported so far (32). However, the targets’ *in vivo* relevance for PV treatment remains elusive (33).

Besides desmosome stabilization, acantholysis is controlled *in vitro* or even *in vivo* through DSG3 expression. The single nucleotide polymorphism rs17315309*G is a risk variant for PV. It resides in the promoter of the transcription factor *ST18* and increases its expression, thereby down-regulating *DSG3* transcription (3). Glucocorticoids are known to reduce blister formation in patients. Mao *et al.* showed how hydrocortisone- or rapamycin-treated primary human keratinocytes down-regulate *STAT3*. This caused an increased expression of *DSG3* with increased numbers and longer desmosomes and resistance to PV IgG–induced loss of cell adhesion (34). An unbiased study that combined whole genome DSG3 knockout and promoter screening comprehensively identified potential modulators of DSG3 transcription impacting PV IgG– induced acantholysis. The study identified and validated KLF5 as a novel regulator of DSG3 expression that can protect against PV IgG-inducing autoantibodies (35).

Further involvement of apoptosis and FasL in PV AAb signaling was postulated (36). Inhibition of FasL reduced fragmentation *in vitro* and in a HSOC model for PV (13).

Here, we address the long-standing question (i) whether PV AAbs directly cause downstream cellular signaling events that lead to cell-wide and long-lasting transcriptome and/or proteome changes modulating cell proliferation or apoptosis, or (ii) whether the observed effects are putative secondary due to disturbed desmosome homeostasis and the ensuing reduction of adhesion. Therefore, we performed a longitudinal transcriptome and proteome response in 2D (37) and 3D (38,39) *in vitro* models of PF and PV up to 48 hours after stimulation and evaluated signaling with primary and secondary effects concerning the observed phenotypes of fragmentation and split formation.

## Methods

### 2D NHEK cell culture

Normal primary human epidermal keratinocytes (NHEK) from three juvenile donors (denoted as donors X, Y, Z) (#HPEKs, lot nos. ES1503029, CP1911101, MC1706096) were purchased from CELLnTEC (Bern, Switzerland) at passage 2. The cells were expanded to passage 4 and cryopreserved for later use at a density of 1×10^6^ cells per vial using CnT-CRYO Defined medium (CELLnTEC, #CnT-CRYO-50). For experiments, HPEKs were cultured starting at passage 4 by seeding at an initial density of 4×10^4^ cells/cm² in CnT-07 medium (CELLnTEC, #CnT-07) at 37°C and 5% CO₂. Once the cells reached 80% confluency, they were detached from the culture dish using Accutase Cell Detachment Solution (CELLnTEC, #CnT-Accutase-100) and seeded for experiments. For experiments, cells were seeded at passage 5 at a density of 4×10^4^ cells/cm². Upon reaching 85% confluency, calcium concentration was increased to 1.2 mM using CaCl₂ (Merck, #102382) to stabilize cell-cell adhesion and synchronize differentiation. Six hours after increasing the calcium concentration, cells were challenged with 50 µg/mL PX43 or hIgG (control antibody) and harvested after 5, 10, 24, and 48 hours. RNA was isolated following the manufacturer’s protocol for the RNeasy Mini Kit (Qiagen, #74104), and residual DNA was removed using the RNase-Free DNase Set (Qiagen, #79254). RNA quality was assessed using the NanoDrop™ 2000 spectrophotometer (Thermo Fisher Scientific, #ND-2000), and samples were stored at −20°C until sequencing.

For protein extraction, the same cell culture protocol was followed. Proteins were collected at 1, 5, and 10 hours post scFv or antibody stimulation. Lysis was performed using a Triton X-100 lysis buffer containing 1% Triton X-100 (Sigma-Aldrich, #T8787), 150 mM NaCl (Merck, #106404), 20 mM Tris-HCl (pH 7.5) (Merck, #100317, #108382), 15 mM Na₃VO₄ (Thermo Scientific

Chemicals, #205330500), 15 mM β-glycerophosphate (Sigma-Aldrich, #35675), 15 mM NaF (Fisher Chemical, #S-3880-50), 1.5 mM PMSF (Sigma-Aldrich, #P7626), and cOmplete™ EDTA-free Protease Inhibitor Cocktail (Roche, #COEDTAF-RO). Protein lysates were centrifuged at 10,000 rpm, 4 °C for 10 minutes to separate the Triton X-100-soluble (non-keratin bound) and Triton X-100-insoluble (keratin bound) fractions. The Triton X-100-soluble fraction was used for shotgun proteomics. All experiments were performed with three biological replicates.

#### Keratinocyte dissociation assay (KDA)

The KDA was performed on NHEKs with low and high CaCl_2_ CnT-07 medium with supplements. 74,500 cells were seeded into the wells of a 24-well plate and grown to 100 % confluency. Once this was achieved, the calcium concentration was increased to 1.2 mM for 6 hours. PX43 and hIgG were added to the wells in concentrations of 1, 10, 20, 50, and 100 µg/mL in triplicates. After 24 hours the medium was aspirated, 400 µL of 2.5 U/mL dispase II (Stem Cell, #07913) were added and incubated for 40 minutes under standard cell culture conditions. The cell sheets were pipetted up and down ten times to induce mechanical stress. Previously, the pipet tips were coated with 1 % BSA for at least 30 minutes. Fixation of the fragments was done by the addition of Roti Histofix® (Roth, #P087.3) and stained with 0.1 % crystal violet (Sigma Aldrich, # V5265). The pictures were taken 24 hours later, and the number of fragments was assessed.

### 2D HaCaT cell culture

Plasma samples were obtained from endemic PF patients (n = 4; female, 21 [min]-51 [max] age) and endemic controls (n =3; 2/3 F, 1/3 M, 66 [min]-74 [max] age; no history of autoimmune diseases) from PF endemic areas in Central-West of Brazil, under approval of Brazilian National Committee for Ethics in Research (CONEP protocol CAAE 02727412.4.0000.0096, approval 505.988). DSG1 and DSG3 was quantified in all samples by ELISA. Clinical and demographic data are described in Table SX. IgG antibodies were purified from plasma samples using Pierce™ Protein A IgG Purification (Thermo Fisher Scientific, Waltham, MA, USA), quantified by Bradford Protein assay (Thermo Fisher) and stored at −80 °C until use.

Human immortalized HaCaT cells were grown in DMEM (Gibco, Waltham, MA) supplemented with 10% FBS, at 37 °C and 5% CO_2_. After reaching 80% confluency, HaCaT cells were stimulated in triplicates with 0.5 mg/mL of IgG antibodies pooled from endemic controls (AV1-AV3) and endemic PF (AV4-AV6) for 12 hours. After treatment, long RNAs (mRNAs and lncRNAs) were isolated using the MiRvana miRNA isolation kit (Ambion). Contaminant DNA was removed from RNA samples using the DNase Turbo kit (Thermofisher), according to manufacturer’s instructions. RNA samples were evaluated according to quality and concentration with Bioanalyzer 2100 (Agilent), where they showed RIN > 8.5. Then, 900ng of each RNA sample proceeded to rRNA depletion using the Low Input RiboMinus Eukaryote System v2 kit (Ambion), following manufacturer’s protocol. The final RNA proceeded to cDNA libraries (Ion Total-RNA Seq Kit v2) and total RNA sequencing was performed using the Ion Proton (Ion Torrent, Life Technologies).

### Human Skin Organ Culture model for PV

The Ethical Committee of the Medical Faculty of the University of Lübeck reviewed all experiments with human samples (reference numbers: 12-178 and 06-109), which were performed following the Declaration of Helsinki. All patients gave their written informed consent for the use of the serum or skin for research purposes.

The protocol for the HSOC model has been described in detail elsewhere (39). In brief, human skin was cut into 1cm^2^ sections and stored in a sterile Petri dish containing William’s E medium (BIOSell) with 100 Units (U)/mL penicillin, 100 µg/mL streptomycin and 0.025 mg/mL amphotericin B (all Gibco) on ice until further use (the skin was used within 24 hours after surgery). Prewarmed defined William’s E-medium with 2 mM L-Glutamine (Biochrom), 100 U/mL penicillin, 100 µg/mL streptomycin (Gibco), 0.01 µg/mL hydrocortisone (Sigma) and 1 µg/mL human recombinant insulin (Sigma) was added to the wells of a transwell cell culture insert plate (Sigma-Aldrich) to a height that created an air-liquid interface between the epidermis and the air.

This experiment was performed in five independent skin organ cultures. Human IgG or DPBS served as negative controls to the stimulation with antibody phage display-derived human bispecific monoclonal single-chain variable fragment (scFv) against DSG3 and DSG1 (40,41). Allocation of skin specimens to the different groups was randomly performed. Afterward, the transwell plates were maintained in the incubator for 24 hours. Next, the skin samples were harvested and cut in half with a sterile scalpel. One-half of the skin was fixed in 4% Histofix solution for subsequent paraffin embedding and haematoxylin– eosin (H&E) staining. The other half was embedded in Tissue-Tek Optimal Cutting Temperature (OCT) compound (Sakura Finetek, Staufen im Breisgau, Germany) for direct immunofluorescence staining for the binding of the scFv. For each HSOC specimen, nonoverlapping pictures of one H&E-stained section were taken over the total length of the skin piece (Keyence). Using ImageJ, the percentage of epidermal split formation was quantified by an investigator unaware of the applied treatments. Image-based quantification of split formation in the epidermis of the HSOC using 21-60 visual fields was used to quantify the efficacy of the AAbs and inhibitors.

Inhibition of split formation was done as previously described (12). As a negative control, human IgG or DPBS was used, and PX43 as a positive control. For each HSOC, at least two skin specimens were required as positive control or NC, as well as three skin pieces for testing each compound at different concentrations. First, only the inhibitors were injected into the dermis of the skin pieces (volume: 50 μl) before placing the pieces from each well on a transwell plate. Positive control and NC were injected with DPBS. The transwell plate with the prepared skin was placed in a humidified incubator (37°C, 5% CO2) for 2 hours. Subsequently, the same amount of inhibitor per skin piece was injected together with the scFv.

For the time course experiment, skin pieces were injected with either 85 µg of PX43, 85 µg hIgG, 34 µg AK23, or 34 µg of murine IgG (mIgG). One skin piece was prepared for RNA isolation and fixation in Histofix® shortly before injections took place (T0). The injected skin pieces were cultured for 5, 10, and 24 hours and prepared after the respective incubation times for RNA isolation and fixation in Histofix® with subsequent HE staining.

### RNA isolation, sequencing, and analysis

The isolation of basal keratinocytes for RNA isolation and sequencing was done with the help of the Olympus stereomicroscope by microdissection. For this, a Petri dish was filled with dry ice and the lid was placed on top. On the lid, two glass slides were placed and let to cool down. Forceps, scalpels, and razor blades were kept on dry ice, too. Between the different skin pieces, everything was cleaned with RNase Away. The skin samples for microdissection were kept on dry ice throughout the procedure, as well as the isolated basal layers.

The skin piece was placed on one of the cooled glass slides with the epidermis facing up and cut into 0.5 to 1 mm thin slices with a cooled razor blade; the skin piece was held by cooled forceps and the slices pulled away from the razor blade with a cooled scalpel. Then, one skin slice at a time was placed on the second glass slide, and as much as possible of the epidermis was removed. Then, the dermis was cut off, and the remaining basal layer was transferred with the help of the cooled forceps into an empty Eppendorf tube on dry ice. This procedure was repeated until all basal keratinocytes were microdissected from the skin sample.

After microdissection of the basal keratinocytes, the samples were kept on dry ice until further processing. The releasing (RL), high solution (HS), and low solution (LS) buffers were prepared according to the instructions for the innuPREP RNA Mini Kit 2.0. Once a set of samples was prepared, 200 µL of RL buffer was added to each sample, and the keratinocytes were transferred with the buffer into the innuSPEED Lysis Tube P. The homogenization was done with the help of the Speed Mill for once 20 seconds (sec). The isolation of the RNA was performed according to the instructions of innuPREP RNA Mini Kit 2.0. For quality control, 1 µL of the RNA was measured via the Nanodrop, and the remaining RNA was stored at −80 °C until being sent for RNA sequencing.

The Illumina TruSeq Stranded Total RNA Library Prep Kit with RiboZero Plus rRNA depletion was used for the library preparation. The sequencing was performed on an Illumina Novaseq6000 with 2x 150 base pair read length, and for analysis, the data was de-multiplexed at the NGSP. Raw sequencing reads in FASTQ format were mapped to the human reference transcriptome (GRCh38) using kallisto (42). On average 75.2*10^6^ reads per sample were pseudoaligned to the Ensembl human cDNA sequences (Version 105), resulting in 14.5*10^6^ unique reads. TPM values were imported and converted to gene-level expression via tximport (43). The differential gene expression was quantified by a likelihood ratio test as implemented in DESeq2 (44), considering stimulus and batch effects as the full and reduced models, respectively. Fold changes were corrected by the apeglm shrinkage estimator (45) to reduce noise and preserve large differences.

### Protein isolation

For protein isolation, the microdissected basal keratinocytes were placed into 2 mL of lysis buffer containing 1 % Triton X-100 (Sigma Aldrich, #X100-500ML), 150 mM NaCl (Fisher Bioreagents, #BP358-10), 20 mM Tris-HCl (Invitrogen, #15567-027), 15 mM Sodium-Orthovanadate (Sigma Aldrich, #450243), 15 mM β-Glycerophosphate (Cayman Chemical Company, #14405), 15 mM Sodium fluoride (Roth, #2618.1), 1.5 mM Phenylmethylsulphonyl fluoride (ThermoScientific, #36978), 1 x cOmplete™ EDTA-free Protease Inhibitor Cocktail (Roche, #COEDTAF-RO) in Aqua dest, vortexed briefly and kept on ice. Tissue disruptor (Ultra Tarax T25) and tissue disruptor attachments (21-30750H from Omni TH biolab products) were used to disrupt the basal keratinocytes. After disruption, the samples are kept on ice for 30 minutes and vortexed once after 15 minutes. Once the incubation time is over, the samples are centrifugated at 200 g for 1 minute at 4 °C. The supernatant is transferred into a new, low-protein binding tube, vortexed, divided into five tubes, and snap-frozen with liquid nitrogen. The pellet is washed with 1 mL lysis buffer and centrifugated at 15,000 g for 10 minutes at 4 °C. Finally, the supernatant is discarded, and the pellet is snap frozen with liquid nitrogen. All samples were stored at −80 °C until quality and quantity control was performed.

### Shotgun proteomics

Proteomic intensity data were acquired from the core facility at the University of Bern. Proteins lysed in TX-100 buffer from PX43/hIgG-treated 2D NHEK and 3D HSOC were precipitated with 5 volumes of cold acetone overnight at −20°C. Proteins were pelleted by centrifugation at 16’000g for 10min at 4°C, and the acetone supernatant was discarded. Pellets were dried in ambient air for 15 minutes and stored at −20°C until use. Proteins were re-dissolved in 100μL 8M urea/50mM Tris-HCl pH 8, and protein content was determined by BCA assay after 1:10 (v/v) dilution with water. Proteins were reduced and alkylated as described elsewhere (46). Urea was then diluted to 1.6M by the addition of 20mM Tris-HCl pH 8.0 / 2mM calcium dichloride before digestion of the proteins with trypsin at a 1:100 ratio for two hours at 37°C followed by the addition of an additional trypsin aliquot for overnight digestion at room temperature. Digestions were stopped by adding 1/20-volume of 20% (v/v) tri-fluoroacetic acid (TFA, Fluka). The digests underwent nano-liquid chromatography analysis using a Nano Elute2 (Bruker Daltonics) connected to a timsTOF HT (Bruker Daltonics, Bremen, Germany), via a CaptiveSpray source (Bruker, Bremen, Germany). The analysis was conducted with an endplate offset of 500 V, a drying temperature of 200 °C, and with the capillary voltage fixed at 1.6 kV. For the chromatographic separation, a volume of 5µL corresponding to 500ng protein digest was loaded onto a pre-column (C18 PepMap 100, 5µm, 100A, 300µm i.d. x 5mm length, ThermoFisher) and subsequently eluted in back flush mode onto a PepSep C18 10-cm column 100 x 0.075 mm, 1.9 μm) by applying a 30-minute gradient of 5% acetonitrile to 30% in water / 0.1% formic acid, at a flow rate of 400 nl/min.

The timsTOF HT was operated in DDA or DIA mode using the Parallel Acquisition SErial Fragmentation (PASEF) mode. For the DDA method, the mass range was set between 100 and 1700 m/z, with 8 PASEF scans between 0.75 and 1.35 V s/cm^2^. The accumulation and ramp time were set to 100 ms. Fragmentation was triggered at 15,000 arbitrary units (au), and peptides (up to charge 5) were fragmented using collision-induced dissociation with a spread between 20 and 75 eV. The dia-PASEF acquisition method was set with 47 isolation windows of 26 m/z width, including an overlap of 1m/z. Isolation windows were associated with ion mobility range between 0.7 to 1.45V s/cm^2^. TIMS accumulation and separation were both set at 100 ms. The DDA data were used to produce a spectral library with fragpipe (47) version 20.0 with the following parameters: database was SwissProt Homo Sapiens (version 2023_04), to which common contaminants were added; precursor and fragment mass tolerances were set to ±20 ppm and ±0.05 Da, respectively; protein digestion was set to trypsin, with a maximum of 3 missed cleavages; variable modifications allowed were oxidation on methionine and protein N-terminal acetylation; carbamidomethylation of cysteines was given as fixed modification; the minimum number of matched fragments was set to 5; validation was done with Percolator (MSBooster enabled); filtering was done at 1% FDR for protein level. DIA data was analyzed by DIA-NN 1.1.8.2 and integrated into the fragpipe suit using the spectral library.

Potential contaminants were then removed for further analysis. Missing DIA protein intensity values were imputed in the following manner: if there was at most 1 detection in a replicate group, then the remaining missing values were imputed by a random draw from a Gaussian distribution of width 0.3 x sample standard deviation and shifted left from the sample mean mu by 2.5 x sample standard deviation; all other missing values were replaced by the Maximum Likelihood Estimation method (48) for differential expression tests using the paired moderated t-test (49). For multiple test corrections, the R tool fdrtool (50) was applied. Significance criteria using 20 imputation cycles were applied as described in Ref. (51) (paragraph 4.9), imposing a minimum log_2_ fold change of 1 in absolute value and a maximum adjusted p-value of 0.05 (the latter value reachable at asymptotically high log_2_ fold changes). After acquisition, the protein intensity values were subjected to logarithmic transformation (base 2) to normalize the data distribution. Differential protein abundance was analyzed using limma (52), which employs generalized linear modeling coupled with empirical Bayes methods to enhance statistical inference. This approach facilitates the robust identification of proteins that exhibit significant changes in abundance across experimental conditions.

#### Bioinformatics Analysis

We calculated the genes/pathways that were differently regulated over time relative to the controls using an F-test as implemented in limma. This is equivalent to a one-way ANOVA for each gene/pathway where the residual mean squares are moderated between genes, additionally. The contrasts for the time series are designed as 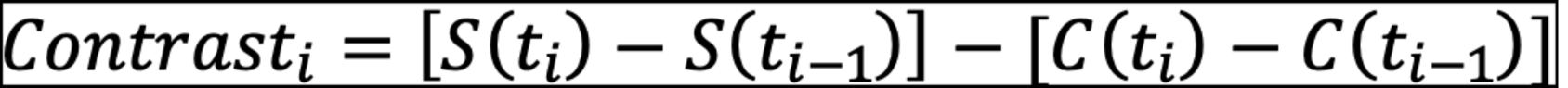, where t, S, and C denote the time points, the respective AAb stimuli, and mIgG/hIgG controls, respectively. Gene set enrichment analysis (GSEA) of REACTOME pathways (53,54) was done by R package gage (55) using all the genes and their log_2_ fold change values. Gene set variation analysis (GSVA) (56) was applied using scaled TPM values per sample.

The significance of the correlation analyses on the fold changes was done using bootstrapping. Symbols were randomly permuted, and the Pearson correlation was recalculated 10.000 times. A skew-normal distribution (57) was fitted to the ensuing distribution of correlation values, from which the respective p-value of the non-permuted fold changes was determined.

Principal Component Analysis (PCA) was calculated using the PCAtools (R package version 2.16.0) library from Bioconductor using the 10% most variable log2-transformed TPM values across all samples. Ellipsoid grouping of data points was based on 95% confidence intervals.

### Data availability

The transcriptome data is accessible from GEO under the ID GSE285010.

## Results

### Experimental rationale for unbiased analysis of keratinocyte response to pemphigus vulgaris autoantibodies

To investigate whether PV and PF AAbs actively trigger long-term, cell-wide signaling pathways that induce a transcriptome response and alter the keratinocyte phenotype or whether they simply prime acantholysis by weakening cell-cell adhesion, we quantified the transcriptomic response to PF-IgG. Additionally, we analyzed the response to DSG1- and DSG3-targeting AAbs in a 2D cell culture model and a HSOC model for PV over 24 hours.

Our experimental design is based on the hypothesis that if IgG and AAbs initiate cellular signaling, they will trigger significant downstream changes in gene expression mediated by transcription factors, potentially altering cellular phenotypes. This hypothesis predicts that transcriptomic changes within 24 hours will correspond to phenotypic shifts, such as keratinocyte migration (58), proliferation (59), or differentiation (60), which are markers of these changes.

Previous studies have demonstrated that keratinocytes respond to stimuli such as hepatocyte growth factor (HGF), fibroblast growth factor (FGF), and interleukin-1α (IL-1α) with precise and essential transcriptomic adaptations, emphasizing the validity of our approach (58,61).

### Autoantibody stimulation yields neither a significant transcriptome response nor a change in proteome abundance in a 2D pemphigus model

In a 2D cell culture, we recorded the NHEK transcriptome response at 5, 10, 24, and 48 hours and the proteome response at 1, 5, 10 hours after treatment with PX43 and control human IgG (hIgG) (**Fig. 1A, left)**. A keratinocyte dissociation assay (KDA) confirmed the efficacy of the PX43 treatment over 24 hours, inducing a significantly higher number of cell fragments compared to hIgG after applying mechanical stress (**Fig. 1A, right**). We compared the NHEK transcriptomes across time, donors, and experimental conditions using a principal component analysis (PCA) calculated from the gene-summarized transcripts per million (TPM) expression using the 10% most variable genes. The PCA showed the strongest sample separation with time, indicative of keratinocyte differentiation, followed by donor-specific gene expression. A trend toward sample separation between PX43 treatment and hIgG controls was not observable before 48 hours (**Fig. 1B**). The result was confirmed in a Gene Set Variation Analysis (GSVA) on the HALLMARK gene sets. Over time we found a down-regulation of proliferation-related processes, such as G2M checkpoints, Myc and E2F targets, and a late up-regulation of proliferation-stop and differentiation-related processes such as Myogenesis, P53, and KRAS signaling, irrespective of the treatment (**Fig. 1C**). The Pearson correlation of all pathway activity activities across batch group and time between PX43 and hIgG was near identical (r=0.97).

**Figure 1:**
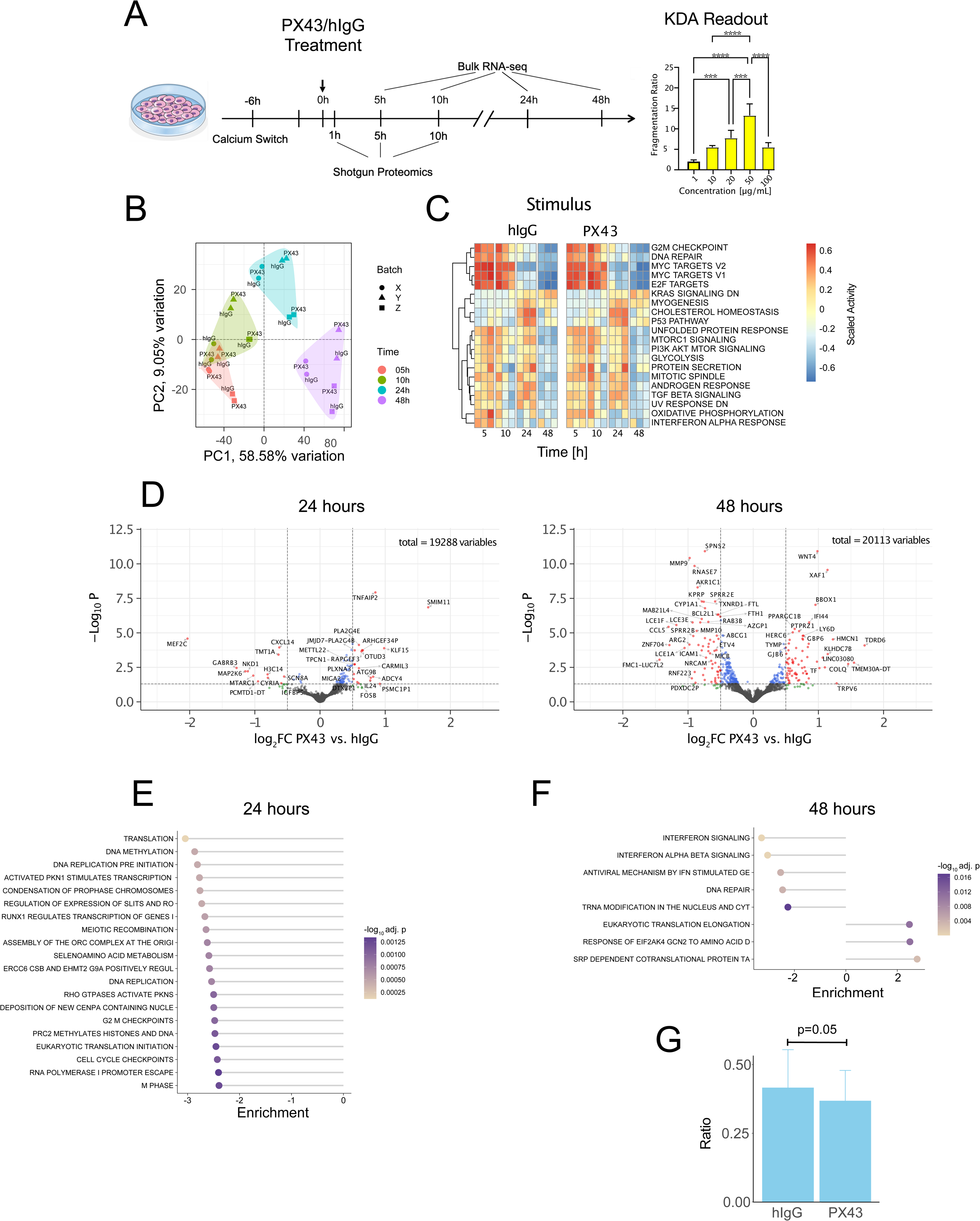
**(A)** Experimental design of the NHEK culture and transcriptome and proteome measurements. Barplots on the right depict the fragment ratio of PX43 vs. hIgG treated cells (n=5) with increasing PX43 concentration after performing a keratinocyte dissociation assay. 3(4) stars denote a p-value significant of 0.001 (0.0001). (B) Principal component analysis (PCA) from NHEK transcriptomes from three donors X, Y, Z after stimulation with either PX43 or hIgG. The PCA has been calculated from the 10% most variable transcripts based on the TPM values. The colored backgrounds group the samples by time point to guide the eye. (C) Heatmaps depicting the activity scores from the MSigDB HALLMARK gene sets based on a Gene Set Variation analysis (GSVA). (D) Volcano plots of the differentially regulated genes between PX43 and hIgG stimulation at 24 hours (left) and 48 hours (right). Transcripts in red are considered significant at an adjusted p-value < 0.05 and a log_2_ fold change > 0.5. (E) Enrichment plot of the 20 most up- and down-regulated REACTOME pathways between PX43 and hIgG at 24 hours from a Gene Set Enrichment Analysis (GSEA) using gage, considering paired samples. The length and direction of the horizontal bars correspond to the mean statistics of GSEA analysis, while the colors correspond to the adjusted p-value. A positive (negative) enrichment corresponds to gene sets up-regulated (down-regulated) in AK23. (F) Same as in (E), but comparing PX43 versus hIgG at 48 hours. (G) Ratio of differentiated to undifferentiated keratinocytes in the 2D NHEK culture 48 hours after the respective stimulation. Error bars denote the standard deviation. p-values stem from a paired t-test on the ratios per batch.

With the transcriptome responses of the PX43 and the hIgG treatment groups being this similar, a time series analysis of the PX43 stimuli relative to hIgG controls yielded neither differentially regulated genes nor HALLMARK or REACTOME pathways, if based on the respective limma or GSVA activity scores (moderated F-test < 0.05). The result is unsurprising when considering differentially regulated genes per time point. Accounting for within-donor correlations (limma duplicate correlation analysis), few genes were regulated (**cf.** volcano plot for PX43 vs. hIgG in **Fig. 1D**; adjusted p-value < 0.05, |log2 fold change| > 0.5) and no gene was consistently differentially regulated over time (**Suppl. Fig. 1A**, |log2FC| > 0.5; adj. p-value < 0.05). Likewise, pathway regulation differed greatly per time point based on a paired gene set enrichment analysis (GSEA). At 5 and 10 hours after PX43 treatment, we found an upregulation of interferon signaling and DNA methylation **(Suppl. Figs. 1B and C, Supplementary Table 1),** while at 24 hours translation, DNA methylation, and replication pathways were downregulated (**Fig. 2E**). Contrary to this, interferon signaling, and DNA repair, as well as translation elongation, were significantly upregulated at 48 hours (**Fig. 2F**). Thus, PX43 treatment did not elicit a long-lasting, consistent transcriptome response over time relative to hIgG control.

**Figure 2:**
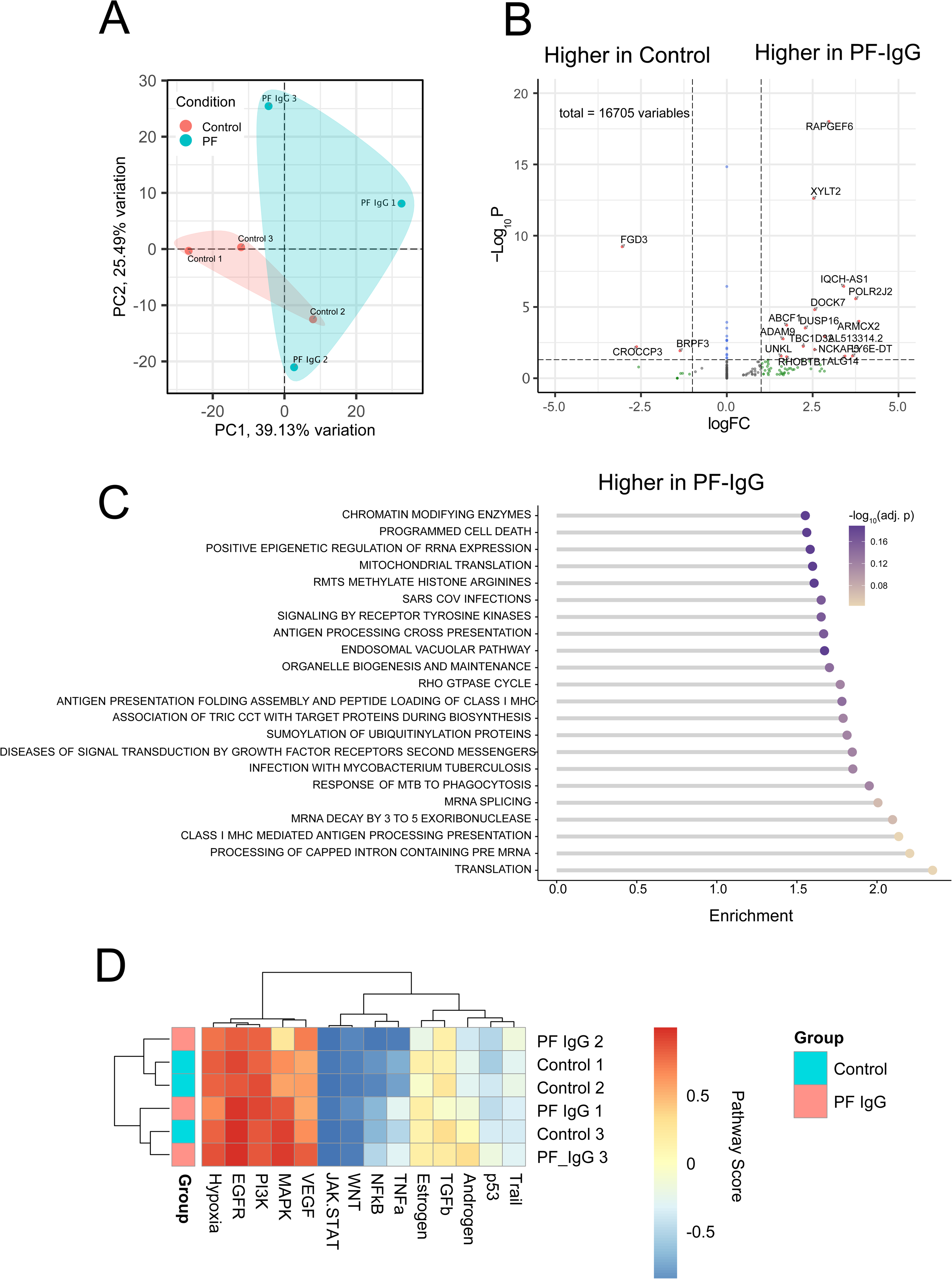
Transcriptome response in HaCaT cells after endemic PF-IgG stimulation. **(A)** Principal component analysis of HaCaT transcriptomes from three donors after stimulation with either endemic PF-IgG or control IgG. PCA has been calculated from the 10% most variable transcripts. The colored backgrounds group the samples by time point to guide the eye. **(B)** Volcano plot of differentially regulated genes between the endemic PF-IgG and control samples. Gene regulation is considered significant above a |log2FC| > 1 and adjusted p-value < 0.01. **(C)** GSEA analysis of PF-IgG vs. IgG. Enrichment denotes the GSEA statistics, while the p-value significance is color-coded. **(D)** Heatmap depicting the upstream pathway activity according to Progeny, based on the transcriptome TPM values. Clustering was performed using average linkage with Euclidean distance as the distance metric.

To better understand this result, we deconvoluted our bulk RNA-seq data with a single-cell RNA-seq experiment from human epidermis skin (62), to obtain the relative abundance of differentiated and undifferentiated keratinocytes. **Fig. 1G** depicts the ratio of the differentiated to undifferentiated cells in the 2D culture. Interestingly, at 48 hours the ratio was significantly higher after hIgG compared to PX43 treatment, which might stem from the transient downregulation of cell-cycle processes, which might stem from PX43 inducing stress, thereby delaying keratinocyte differentiation.

Lastly, we checked whether a changed protein abundance mirrors the changes in gene expression. We quantified 4,515 proteins using shotgun proteomics across all conditions and time points. A principal component analysis revealed a strong separation with batches (**Suppl. Fig. 2A**) and separation over time for the first two and third principal components (PC2=14.4%; PC3=11.3%, **Suppl. Fig. 2B**). The limma duplicate correlation of 0.29 corroborated the results, indicating a strong donor effect. An analysis for differentially regulated proteins, accounting for the donor effects, revealed 55, 34, and 23 proteins as differentially regulated between PX43 and hIgG at 1, 5, and 10 hours, respectively (adjusted p-value < 0.05, |log2 fold change| > 0.5; **Suppl. Fig. 2C**). A GSVA using HALLMARK gene sets showed E2F processes and interferon alpha response up-regulated, while IL-6 and IL-2 as well as JAK-STAT responses were down-regulated over time, independent of PX43 treatment (**Suppl. Fig. 2D**).

In summary, PX43 neither induced long-lasting cell-wide changes in the transcriptome nor proteome of a 2D NHEK culture under high calcium conditions over time. Donor-specific and keratinocyte differentiation effects had a much stronger effect on the transcriptome and proteome than any change due to AAb stimulation. However, we found a transiently induced, cellular stress response that delayed the transition from proliferation to differentiation relative to the hIgG controls.

### Endemic Pemphigus foliaceus IgG does not induce a significant transcriptome response in a 2D culture of HaCat cells

Having rejected the notion of a long-lasting, concerted transcriptome response in NHEK cells after stimulation with DSG1 and DSG3 AAbs, we next evaluated whether 2D epidermal keratinocytes exposed to endemic PF IgG derived from patient plasma induce a downstream gene response related to split formation, cell detachment, or wounding. For this, we quantified the transcriptome responses to IgG AAbs from the peripheral blood plasma of patients with endemic pemphigus foliaceous (PF) compared to IgG from endemic PF controls. PF IgG targets DSG1 and thus induces sub-epidermal loss of intercellular adhesion. In the 2D *in vitro* assay, HaCaT cells were grown to 80% confluence in culture dishes and stimulated with patient IgG or control for 12 hours. A PCA on the 10% most variable transcripts values indicated a large inter-sample heterogeneity and an incomplete separation of sample groups (**Fig. 2A**). Consequently, there were only a few, namely (17/3) up-/down-regulated genes between the controls and the endemic PF-IgG treated samples (adj. p-value < 0.05, |log_2_ Fold Change| > 0.5, **Fig. 2B**). A Gene Set Enrichment Analysis (GSEA) indicated a significant up-regulation of translation, processing of capped intron and antigen processing presentation in endemic PF-IgG treated cells (**Fig. 2C**), but no significantly down-regulated pathways. Moreover, none of the signaling pathways are related to inflammation, cell detachment, or wounding. In line with this finding, an analysis of putative upstream pathways using Progeny did not reveal any differences between the two groups (**Fig. 2D**) and did not reflect any of the known signaling pathways that are known to be regulated after AAb binding, like P38-MAPK, or the endoplasmatic reticulum (63).

In summary, PF IgG did not induce cell-wide, concerted transcriptional changes in a 2D HaCaT culture. Donor-specific effects were stronger than changes in gene expression after IgG stimulation. Gene regulation did neither reflect a loss of cell adhesion nor was it connected to known protein signaling after PF IgG binding Thus, endemic PF-IgG had little to no effect on gene expression within a period of 12 hours.

### Autoantibody treatment changes keratinocyte transcriptome only when associated with split formation in a human skin organ culture model for pemphigus vulgaris

To find out whether the above findings hold for PV antibodies in a 3D cell culture as well, we next investigated the transcriptome and proteome responses over 24 hours in basal keratinocytes in an HSOC model after AK23 or PX43 AAb treatment. AK23 (64,65), is a DSG3 murine IgG1 antibody, while PX43 targets both DSG1 and DSG3. Due to their origins, we used murine IgGs (mIgG) and human IgGs (hIgG) preparations as controls, respectively. Treatment and control antibodies, as well as inhibitors, were injected close to the dermal-epidermal junction at specified time points (**Fig. 3A**), and cultures were sampled for split formation, transcriptome, and proteome analysis after 24 hours, with additional transcriptome sampling at 5 and 10 hours.

**Figure 3:**
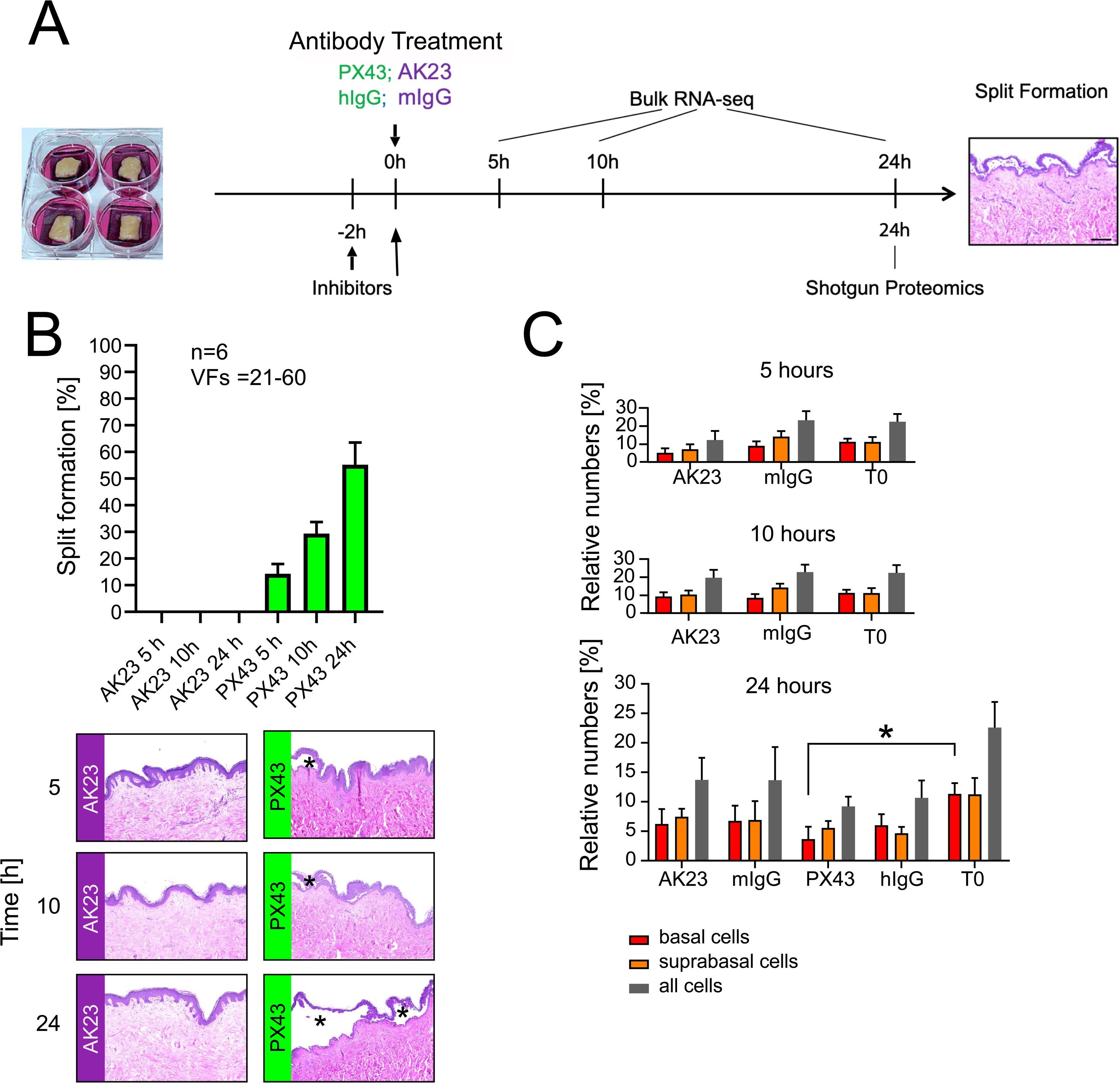
Experimental design and split formation in a 3D human skin organ culture model. **(A)** 3D human skin culture was treated with PX43 or AK23 autoantibodies and respective controls (hIgG, mIgG) by direct application into the dermis close to the basal membrane. RNA or protein from keratinocytes proximal to the basal layer was extracted at [5h, 10h, 24h] and [0h, 24h] after stimulation for bulk RNA-seq or shotgun proteomics, respectively. **(B)** quantification of split formation from 21-69 fields of view from n=6 biological replicates after AK23 or PX43 stimulation. Bottom: representative skin cross-sections at three different time points, black asterisks mark epidermal splits. No split formation was visible after AK23 stimulation. **(C)** Quantification of Ki67 staining in basal, suprabasal, and the combined cell populations before (T_0_), at 5, 10, and 24 hours after AK23/mIgG and PX43/hIgG treatment. No significant differences between time points and conditions were observable, except a reduction in Ki67 between T_0_ and PX43 basal keratinocytes after 24 hours (p-value < 0.05).

Reflecting the phenotype in the patient, split formation was absent for all time points after AK23 stimulation as well as mIgG and hIgG injection (data not shown)(**Fig. 3B**). In contrast, injection with PX43 resulted in increased split formation over time, which was detectable after 5 hours already. The absence of split formation after AK23 stimulation was likely due to targeting DSG3 only, which left compensatory DSG1-mediated cell-cell adhesion unaffected (66,67). Since previous studies on an adult mouse model for PV showed increased epidermal proliferation two hours after AK23 administration (68), we examined whether a similar phenotype could be observed in the HSOC model. However, Ki-67 staining of proliferating basal and suprabasal epidermal keratinocytes showed no significant differences between the AAb-stimulated and control groups (**Fig. 3C**), indicating that neither of the two AAbs increased cell proliferation in this case.

We quantified the transcriptome response to the AAbs and IgG controls by extracting basal keratinocytes from the HSOC model before stimulation (0 h) and at 5, 10, and 24 hours after injection (n=6). A PCA revealed transcriptional changes with time, but also for PX43 compared to hIgG (compare filled squares (PX43) vs. crosses (hIgG) at 24h in **Fig. 4A**). In line with the observed split formation phenotype and the PCA, PX43 induced an increasing number of significantly deregulated genes over time relative to hIgG control (**Fig. 4B** and **Table 1**, adjusted p-value < 0.01, |log2 Fold Change| > 1, cf. **Supplementary Table 2**). In contrast to the 517 up- and down-regulated genes 24 hours after PX43 stimulation (**Fig. 4B**, inset bottom), only 7 genes were significantly deregulated after AK23 treatment relative to mIgG (**Fig. 4B**, inset top). Among the most up-regulated genes after PX43 stimulation, we found interferon (*IFIT3*, *IFI44*), cytokines (*CXCL10*, *CXCL11*), and interleukins (*IL1B*). To identify pathways with significant regulation over time, we conducted a time series analysis on pathway activity scores from a GSVA using REACTOME gene sets. While AK23 stimulation did not produce any temporally regulated pathways, PX43 stimulation led to differential regulation in 120 gene sets (**Fig. 4C**, limma F-test, adjusted p-value < 0.001, **Supplementary Table 4**). Among the most upregulated pathways were those related to inflammation, including interleukin, interferon, NFkB, and TRAIL signaling. It is also worth noting the weaker and transient regulation of inflammation and interferon signaling in samples stimulated with AK23, hIgG, or mIgG at the 5- and 10-hour time points, which subsides by 24 hours. This may result from the physical injection of AAbs/IgG, creating internal pressure at the dermal-epidermal junction.

**Figure 4:**
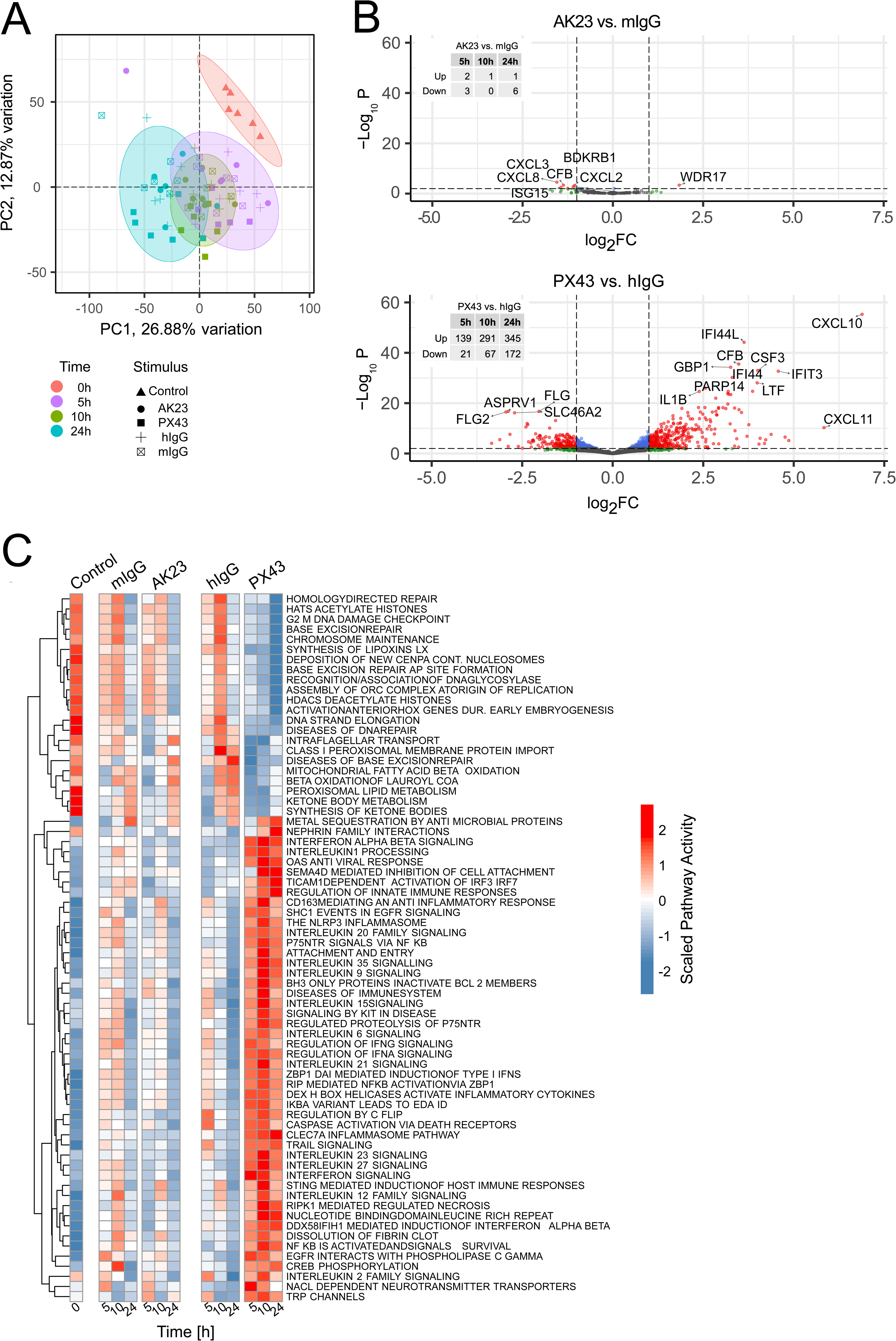
Transcriptome response upon autoantibody stimulation in a 3D human skin organ culture model for pemphigus vulgaris. **(A)** Principal component analysis of transcriptomes from basal keratinocytes extracted at the indicated time points after the respective stimuli. The percent values denote the explained variance per principal components (PCs). Ellipses enclose the data points with respect to time at a 95% confidence interval. **(B)** Volcano plots of the differentially regulated genes at 24 hours for AK23 vs. mIgG (top) and PX43 vs. hIgG (bottom). Genes are marked as significant at an adjusted p-value < 0.01 and a |log2 fold change| > 1. The inset tables provide the respective numbers of differentially up- and down-regulated genes for the different time points. **(C)** Heatmap of the row-wise scaled, averaged pathway activity scores (n=6) from a GSVA analysis showing the significantly regulated REACTOME pathways over time after PX43 stimulation relative to hIgG. Rows were clustered using average linkage with Euclidean distance as the distance metric.

**Table 1:** Number of differentially regulated genes per time point after AK23 or PX43 stimulation relative to the respective mIgG and hIgG controls in the 3D HSOC model. Cutoff thresholds for differential regulation are adjusted p-value < 0.01 and an |log2FC| > 1.

| Time |  | 5h | 10h | 24h |
| --- | --- | --- | --- | --- |
| Donor correlation |  | 0.01 | -0.06 | -0.2 |
| AK23 | Up | 2 | 1 | 1 |
|  | Down | 3 | 0 | 6 |
| PX43 | Up | 139 | 291 | 345 |
|  | Down | 21 | 67 | 172 |

We then identified differentially abundant proteins using a shot-gun proteomics approach (cf. **Supplementary Table 3**), comparing pre-stimulus (T_0_) and 24 hours after PX43 or hIgG treatments (n=5). In total, 3,720 proteins were detected in all samples. A PCA revealed a strong clustering of the 0 and 24-hour time points (**Fig. 5A**), with a considerable donor-specific consensus correlation of 0.53 (limma duplicate correlation analysis). A robust linear regression on the PX43 and hIgG samples identified 199 differentially regulated proteins (101 up- and 98 down-regulated) based on the log_2_-transformed imputed protein intensities (**Fig. 5B**, adjusted p-value < 0.05, cf. **Supplementary Table 3**). The most up-regulated proteins were related to inflammation and the immune system, such as S100A8/9, TNFAIP6, LCN2, and IL6. The differential protein abundance after 24 hours between PX43 and hIgG correlated positively with differential gene expression (r=0.45, p< 2.2*10^-16^, all transcripts and proteins with adjusted p-value < 0.05), confirming the prior results. A GSEA using gage found eight up- and 17 down-regulated pathways between PX43 and hIgG (**Fig. 5C**, adjusted p-value < 0.05), with interferon and antigen presentation being the most up-, and complement activation the most down-regulated gene sets.

**Figure 5:**
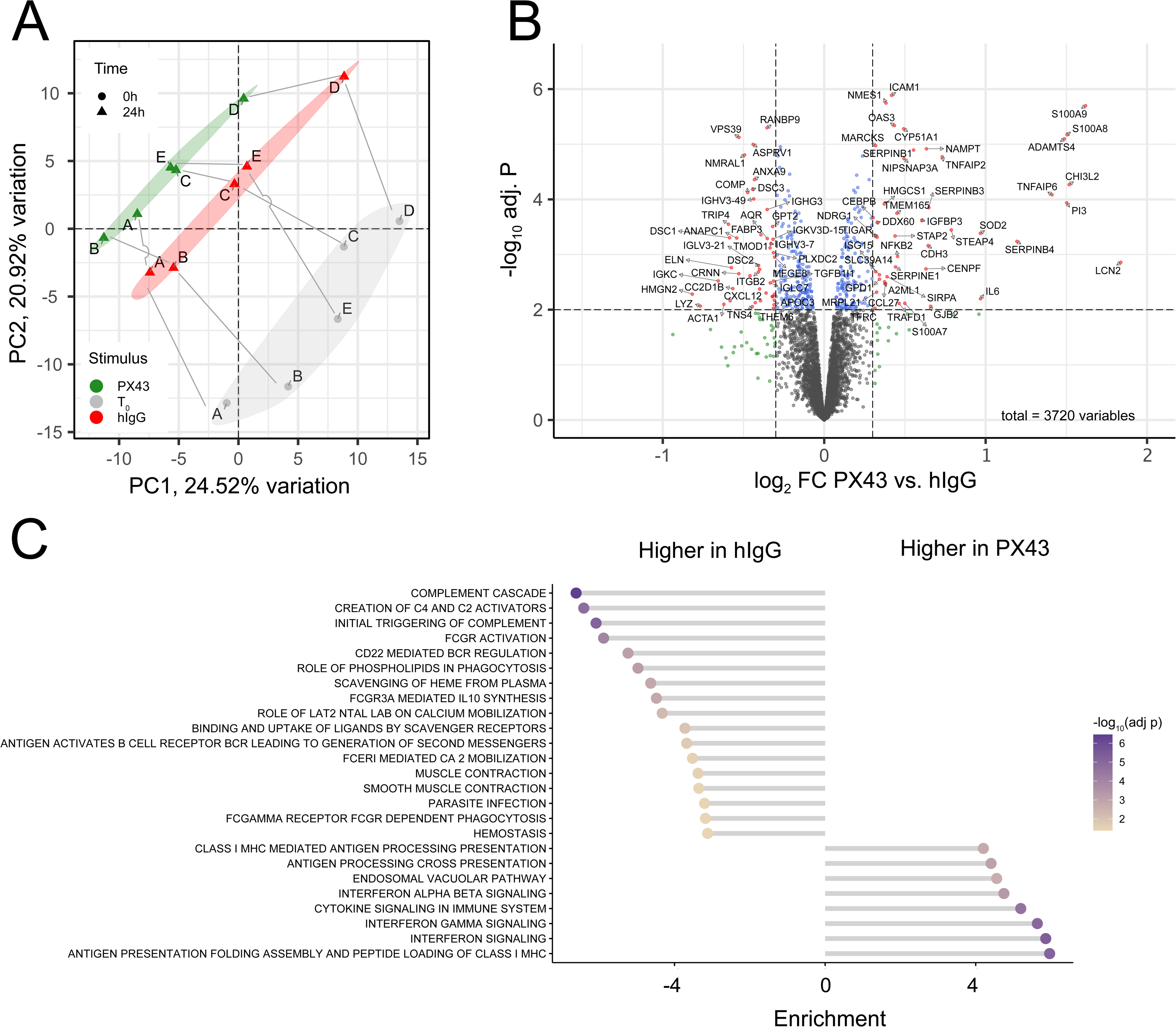
Proteome response analysis after PX43 stimulation. **(A)** Principal component analysis of the 10% most variable proteins before stimulation (T_0_) and after 24 hours of PX43 or hIgG stimulation. Letters mark the biological replicates, and gray lines connect the samples batch-wise to guide the eye. The percent values denote the explained variance per PC. **(B)** Volcano plot of the differentially regulated proteins after a robust linear regression analysis. Significantly regulated proteins (adjusted p-value < 0.01, |log_2_FC| > 0.3) are marked in red and denoted by the respective gene symbol. **(C)** GSEA of the most significantly regulated REACTOME pathways (adjusted p-value < 0.01). The length and direction mark the mean GSEA statistics for up- or down-regulated pathways, using hIgG as a reference. Colors correspond to the adjusted p-value.

In conclusion, we confirmed a long-lasting, cell-wide transcriptome and proteome response associated with the occurrence of split formation, which in turn was linked to inflammation, interferon interleukin activity, and pathogen response. The sole presence of AAbs without visible acantholysis, as in the case of AK23, was insufficient to induce a consistent transcriptome response.

### Differentially regulated genes after PX43 treatment are related to keratinocyte detachment and early wounding and mirror the transcriptome of pemphigus vulgaris skin

From the findings above, we hypothesized that the observed gene response is an indirect result of split formation rather than a direct response to AAb treatment. If the former is true, we expect differential gene regulation after split formation in the HSOC model to relate to cell detachment and early wounding. To validate our hypothesis, we analyzed the correlation between differential gene regulation in PX43 and hIgG with the following transcriptome responses: (i) keratinocyte migration upon hepatocyte growth factor stimulation in a 2D culture (58), (ii) NHEKs differentiating in a 2D culture after switching to high calcium (this study), (iii) NHEKs in a 2D culture after double-paracrine stimulation (GEO ID: GSE271501), (iv) keratinocytes stretched in a microflow device(69), (v) keratinocyte gene response from acute wounding of patient skin (70) and(vi) detached keratinocytes in a wound model (71). For each comparison, we intersected the most significantly regulated genes after PX43 stimulation at T=5h, 10h and 24h (adjusted. p-value < 0.01) with the expressed genes from (i-v) and correlated the log_2_ fold changes relative to the respective controls. As an example, the scatter plot in **Fig. 6A** compares the fold change values after PX43 treatment with the transcriptome response from isolated and trypsinized NHEKs relative to micro-dissected epidermis after 16 hours. Correlating 693 common genes, we found a positive Spearman correlation of ρ=0.50.

**Figure 6:**
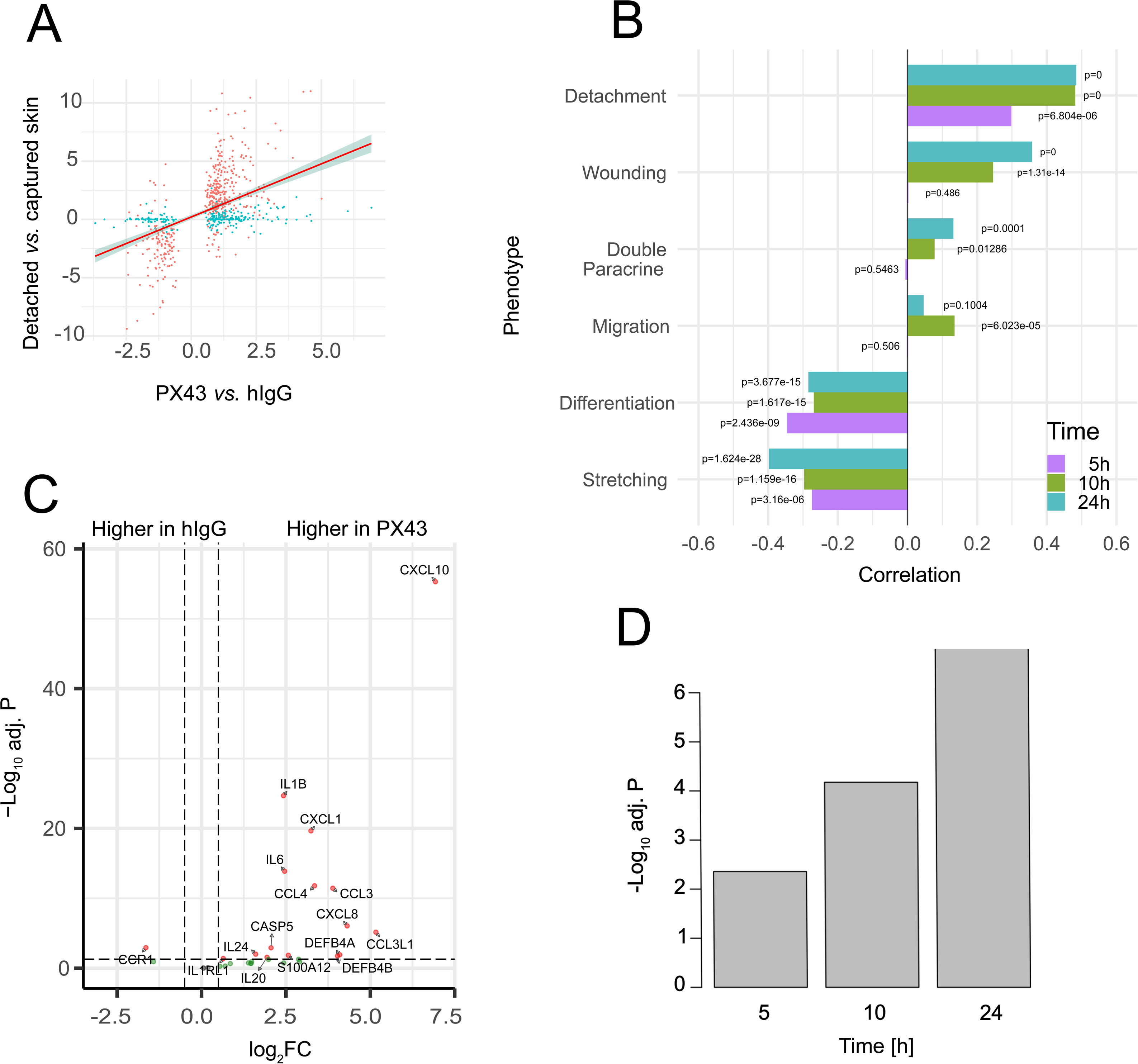
Transcriptome response correlations. **(A)** Scatter plot of log_2_ fold changes between PX43 and hIgG (x-axis) and detached keratinocytes vs. controls (y-axis). The red line denotes a linear fit together with 95% confidence intervals. **(B)** Correlation of the PX43-initiated transcriptome response with keratinocyte gene response to different phenotypes and time. The bar length denotes the Spearman correlation. The associated p-values are noted next to each bar. **(C)** Volcano plot between PX43 and hIgG at 24 hours of 40 genes identified as differentially regulated in PV skin. Significance levels are |log_2_FC| > 1 and adjusted p-value < 0.01. **(D)** Barplot of the adjusted p-value of a GSEA analysis using the 40 genes from (C) as a gene set when comparing PX43 vs. hIgG at 24 hours.

**Figure 7:**
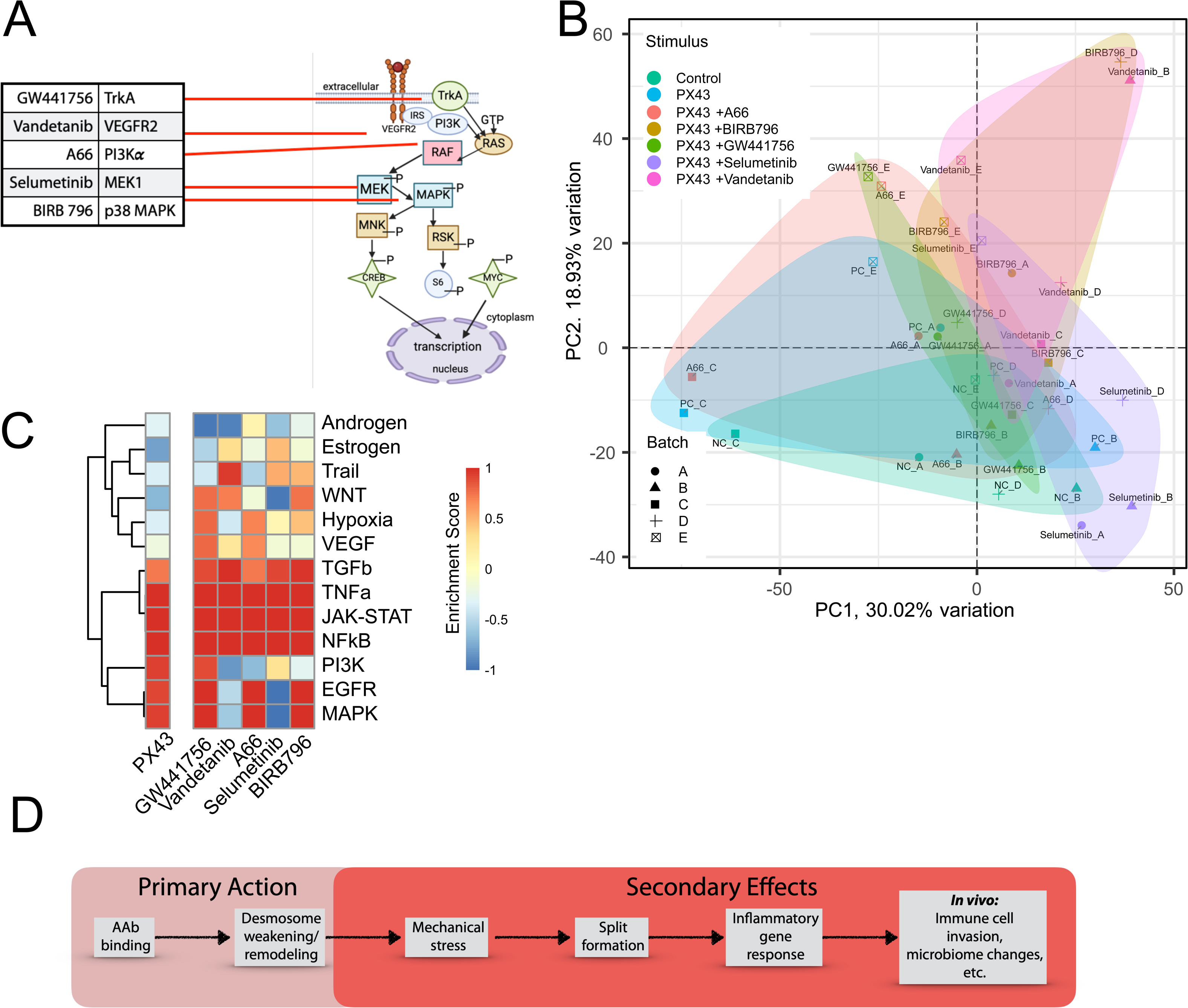
Transcriptome response of an HSOC model to PX43 stimulation and inhibitors of split formation. **(A)** Pathway scheme showing the different inhibitors, the protein targets, and their location in cellular pathways. **(B)** PCA analysis on the gene expression using TPM values on the 10% most variable transcripts. The colored ellipses guide the eye to group the samples according to their stimulus. **(C)** Heatmap of the enrichment scores of upstream pathway activity relative to the control of different stimulus conditions. Rows are hierarchically clustered by complete linkage according to their Euclidean distance. **(D)** Scheme distinguishing between primary and secondary effects of PV-related AAbs.

In passing, we note that the production of AAbs and IgG resulted in background endotoxin levels in PX43, to some extent in mIgG, to a lesser extent in AK23 but not in hIgG preparations. This effect is visible from the fraction of genes regulated under PX43 relative to hIgG but not after cell detachment (**Fig. 6A**, blue dots). A hypergeometric test revealed that these genes are significantly enriched for endotoxin-related genes (p=2.3*10^-76^), thus providing a visual cue to the effect of contamination and cell-detachment effects. To exclude endotoxin-related effects in our results, we recalculated the GSVA-based time series analysis, excluding known genes associated with endotoxin responses (**cf. Supplementary Table 4**). Endotoxin response genes were identified by compiling all differentially regulated genes when comparing endotoxin-contaminated with endotoxin-free mIgG and hIgG over time. This analysis yielded 664, 1,237 and 927 deregulated genes across the three timepoints (p-value < 0.05), for 2,538 unique endotoxin-related genes. When recalculating temporal pathway regulation without these genes, we observed 54 regulated pathways (adj. p-value for F-statistics < 0.001). A Pearson correlation of r=0.85 on the F-statistics between pathway analyses with and without endotoxin-related genes (p-value < 2×10⁻¹⁶) confirmed that endotoxin contamination did not qualitatively impact the gene response under PX43 stimulation.

Having excluded a significant effect of endotoxins on our results, we next determined the significance of the correlation. We applied bootstrapping, generating a skew-normal distribution based on Spearman correlation values obtained by randomly permuting fold change values and recalculating the correlation 10,000 times. **Figure 5B** shows the Spearman correlations and bootstrapping significances across all six phenotypes and three time points. The analysis reveals that PX43-induced effects are significantly correlated with cell detachment and inversely correlated with differentiation and cell stretching, even at early time points. Correlation with acute wounding strengthens over time, whereas correlations with cell migration and paracrine communication appear weaker. This links differential gene regulation to the formation of epidermal splits and the ensuing keratinocyte response as a secondary effect to PX43-induced reduction in cell-cell adhesion, but not the AAbs themselves.

Finally, we assessed whether the gene response observed in the HSOC model for PV aligned with the *in vivo* conditions of patient skin. Holstein *et al.* used bulk RNA-seq to compare skin specimens from PV patients (n=6) with healthy skin and identified 209 significantly deregulated transcripts, selecting 40 differentially regulated genes for further analysis (72), **see Supplementary Table 5**. We found that most of the 40 genes were similarly upregulated following PX43 stimulation and split formation (**Fig. 5C**). A longitudinal GSEA analysis comparing PX43 to hIgG showed increasing significance over time (**Fig. 5D**), indicating a growing similarity in gene regulation between the HSOC model and PV skin.

In summary, the findings support the hypothesis that PX43-induced gene regulation in the HSOC model reflects secondary effects from cell detachment and split formation, aligning progressively with the in vivo gene expression profile of PV skin. These data suggest that observed gene responses are primarily due to mechanical disruption effects rather than the direct effect of AAb stimulation on keratinocytes.

### Inhibitors of split formation yield transcriptome response specific to the keratinocyte detachment and the specific inhibitor

Finding inhibitors of PV-related signaling pathways that stabilize desmosome structures and thereby prevent acantholysis has been the focus of PV research for years (9). Burmester et al. (12) reported on five different kinase inhibitors from a kinase screening library that reduced, but not completely abrogated, split formation in a 2D and a 3D HSOC model (**Fig. 6A**). Here, we asked whether reduced formation in an HSOC model after PX43 stimulation would change the transcriptome response qualitatively or quantitatively, only. For this, we compared the 24-hour transcriptome response of the complete HSOC, i.e., all cells including keratinocytes and fibroblasts, after PX43 stimulation alone and under inhibitor pre-incubation for 2 hours.

A PCA on all samples using the 90% most variable genes showed an incomplete separation of the experimental groups (**Fig. 6B**). An analysis of the causal upstream pathways to each stimulus using Progeny, a linear model for pathway-specific response (73), confirmed the similarities in differential pathway activities. The heatmap in **Fig. 6C** reveals a set of common upstream pathways, namely TGFa/NFkB, TGFb, and JAK-STAT, active in all conditions, irrespective of inhibition. Further pathways, such as EGFR, MAPK, and PI3K, were upregulated after PX43 stimulation but inactivated by Vandetanib (VEGFR2, EGFR and RET inhibitor) and Selumetinib (MEK1 inhibitor), respectively. Thus, despite the added effects of the inhibitors on upstream pathways and split reduction, cell detachment and wounding response pathways remained predominant. The inhibitors did not significantly alter the cellular response, suggesting that the overall transcriptomic profile is primarily driven by the presence or absence of epidermal splits.

## Discussion

Here, we investigated the cell-wide events following the binding of anti-DSG1/3 antibodies and endemic PF-IgG up to 48 hours in 2D and 3D models for PV and PF. Without mechanical insult, split formation was absent in the 2D models after PX43 or endemic PF-IgG treatment. Split formation was likewise absent when incubating the HSOC model with AK23, most likely due to the compensatory DSG1 adhesion in the skin **(Fig. 3A).** Without signs of acantholysis, we found that neither PF-IgG nor AK23 induced significant or concerted long-lasting changes in gene expression or protein abundance in either model. In the case of PX43 treatment, the observed transcriptional changes were not associated with previously known protein signaling pathways in PV or PF, such as EGFR, p38 MAPK, or cAMP signaling, but instead correlated strongly with the occurrence of acantholysis and split formation, resulting in a gene expression profile resembling early wounding and cell detachment. To our knowledge, this is the first comprehensive investigation of transcriptome and proteome changes in the early events of the PV and endemic PF pathophysiology.

These findings have profound implications for understanding the action of pemphigus AAbs. Consistent with existing literature, our data support the notion of PV and PF as desmosome remodeling diseases, where AAbs weaken but do not completely disrupt desmosome-mediated cell-cell adhesion through steric hindrance (22,23,74). Our results are consistent with the clinical Nikolsky’s sign, where rubbing of pemphigus patient’s intact skin induces blistering (75). The results are also consistent with the absence of visible split formation in keratinocyte monolayers. The application of AAbs widens the intercellular space (37,76), yet cells still adhere to each other, even if DSG3 and DSG1 are inactivated through PV-IgG or PX43 (37). Only after additional mechanical force is increased fragmentation observed. While AAb binding causes signaling changes propagating through the desmosome interactome (19,21), these effects appear transient (26) and insufficient to drive large-scale transcriptomic changes. Our findings align with prior studies and systems theory, where transcriptomic alterations are closely tied to cellular phenotypes (58–60,77) and often represent secondary, disease-induced changes rather than primary drivers of disease mechanisms (78). This explains the absence of a cell-wide response in gene expression after AK23/PX43 incubation or stimulation with PF-IgG. If cellular morphology in the respective 2D or 3D *in vitro* models does not change upon AAb treatment and fragmentation or split formation are absent, a well-wide transcriptome response may not happen. However, it cannot be excluded that protein signaling of the desmosome interactome and widening of intercellular spaces induce mild responses of cellular stress, affecting cell proliferation and differentiation, as was evident from the deconvolution analysis that showed reduced cell differentiation in the 2D cell culture (**Fig. 1F)**.

Extending these findings to the *in vivo* situation, we hypothesize that many effects like inflammation (79), immune cell invasion (72), or altered microbiome (80) observed in PV patients’ skin are secondary to split formation rather than a direct consequence of AAb binding **(Fig. 6D)**. Upon binding of the AAbs, desmosomes shrink and decrease in numbers, weakening lateral keratinocyte adhesion. This process is accompanied by protein signaling in the desmosome interactome. Mechanical stress induces split formation and ensuing inflammatory cell response akin to cell detachment and wounding. The latter leads to the PV/PF pathogenesis of immune cell invasion, skin microbiome changes, and further processes.

Of note, our finding contrasts the disease mechanisms in bullous pemphigoid (81). There, autoantibodies (mainly IgG) target BP180 (collagen XVII), a key hemidesmosome protein anchoring keratinocytes to the basement membrane. Autoantibody binding activates the complement system, recruiting immune cells such as neutrophils, eosinophils, and macrophages that release enzymes and pro-inflammatory cytokines, causing tissue damage, and blistering at the epidermal-dermal junction. Notably, keratinocytes can exhibit an inflammatory response to BP180-targeting IgG even without immune cell involvement (82), a mechanism not observed in PV, where DSG3 antibodies primarily weaken adhesion rather than initiate inflammation.

The distinct pathogenesis of PV has significant therapeutic implications. While treatments like corticosteroids and B-cell-depleting therapies (e.g., rituximab) are effective, their side effects, delayed onset, and considerable relapse rates highlight the need for alternative approaches (83,84). Many *in vitro* studies have focused on modulating desmosome-related proteins to reduce fragmentation (9). Pre-treating cells *in vitro* with various inhibitors modulates desmosome turnover and reduces PV-IgG-induced fragmentation (12). However, given that desmosome turnover modulation upon AAb binding is transient and, according to our findings, does not induce long-term transcriptomic changes, the effectiveness of these approaches in clinical settings may be limited. Strengthening desmosome structures themselves, such as through cAMP signaling via apremilast (85) or modulating *DSG3* expression via transcriptional regulators like STAT3, ST18, or KLF5 (3,34,35), represents a promising alternative therapeutic direction. The latter was found. in an unbiased genome-wide screening of DSG3 interactors (35). Interestingly, members of the desmosome interactome did not appear in the promoter screen for *DSG3* regulation. Given our results, one might speculate that desmosome turnover modulation upon AAb binding and transcriptional control of DSG3 expression are disjunct local and cell-wide processes, respectively, emphasizing the need for complementary therapeutic strategies to enhance desmosome stability. In conclusion, our findings confirm steric hindrance including desmosome weakening and internalization as the critical events in pemphigus pathogenesis, highlighting that AAb-induced transcriptional changes are secondary to acantholysis rather than direct triggers of disease mechanisms. These insights open new avenues for therapies targeting keratinocyte adhesion and desmosome stability to combat pemphigus effectively.

## Supplementary Information

**Supplementary Table 1**: Gene set enrichment analysis of differentially regulated REACTOME pathways in a 2D NHEK culture after PX43 vs. hIgG treatment.

**Supplementary Table** 2: Differentially regulated genes in the HSOC model between different conditions and timepoints.

**Supplementary Table 3**: Differentially regulated proteins in the HSOC model between PX43 and hIgG after 24 hours.

**Supplementary Table 4**: Differentially regulated REACTOME pathways in the HSOC model over time using all expressed genes or those unaffected by endotoxins.

**Supplementary Table 5**: Genes used in the comparison from Holstein et al.

## Acknowledgments

HB and SG acknowledge computational support from the OMICS compute cluster at the University of Lübeck. HB, SG, JH, VH, EJM, WVJH, and SR acknowledge funding through the Swiss National Science Foundation grant CRSII5_202301. CMH acknowledges funding from DFG - SFB 1526 Pathomechanisms of Antibody-mediated Autoimmunity, Project-ID 454193335 and funding from the University of Lübeck (CS06-2019). HB acknowledges funding from the Deutsche Forschungsgemeinschaft (DFG, German Research Foundation) under Germany’s Excellence Strategy – EXC 22167-390884018. DM acknowledge funding through the Conselho Nacional de Desenvolvimento Científico e Tecnológico (CNPq protocol: 460668/2014-5). We thank Coordenação de Aperfeiçoamento de Pessoal de Nível Superior [CAPES/PROAP – Finance Code 001] for the scholarships provided to AS-S, VB-BH, VCS, and GAC.

Best regards

---

**Supplementary Figure 1.**
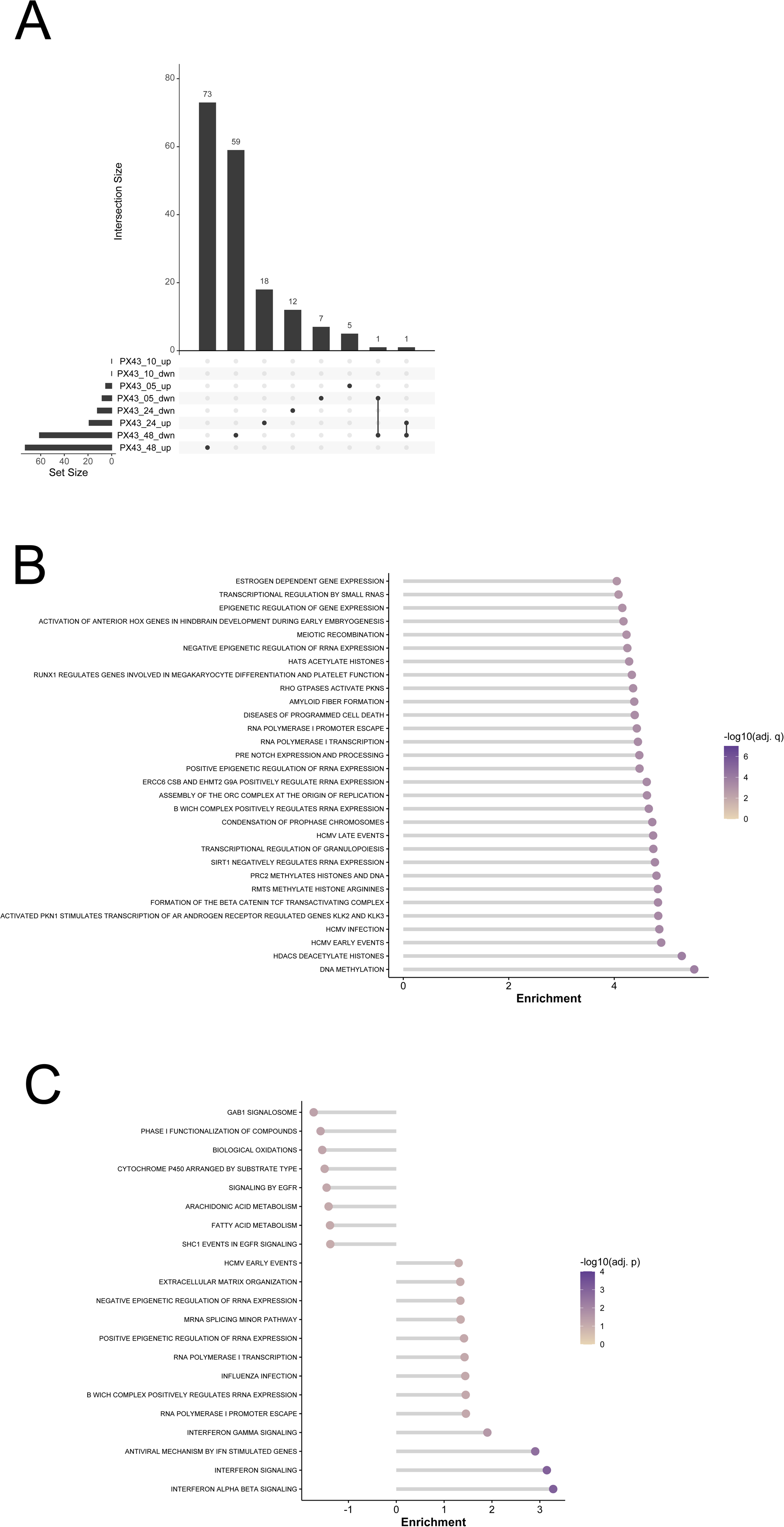

**Supplementary Figure 2.**
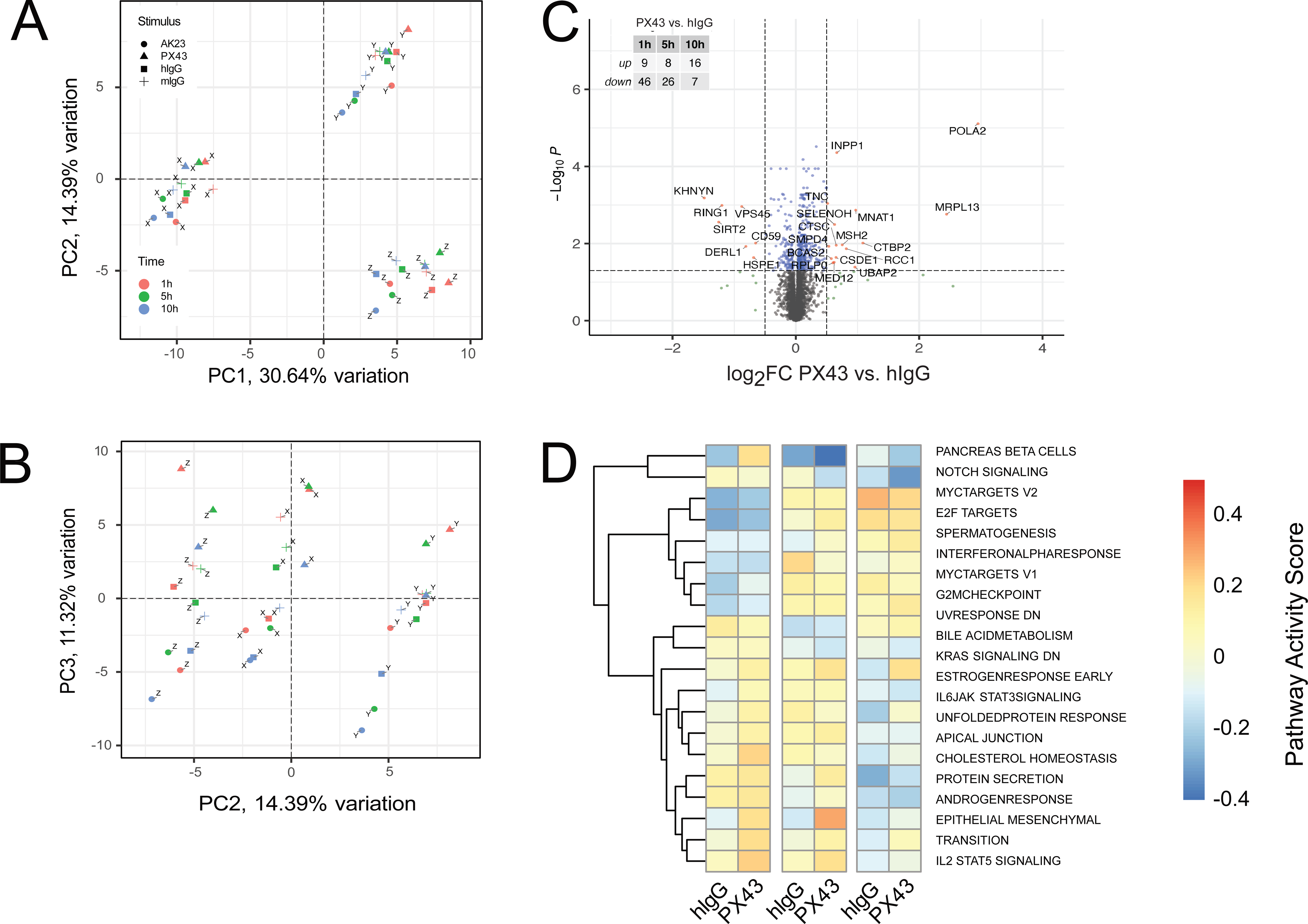

